# Prefrontal cortex neural compensation during an operant devaluation task

**DOI:** 10.1101/2021.08.18.456813

**Authors:** Hayley Fisher, Hongyu Lin, Jensen May, Caitlin McLean, Charles L. Pickens

## Abstract

Deficits in goal-directed action are reported in multiple neuropsychiatric conditions, including schizophrenia. However, dysfunction is not always apparent in early stages of schizophrenia, possibly due to neural compensation. We designed a novel devaluation task in which goal-directed action could be guided by stimulus-outcome (S-O) [presumably orbitofrontal cortex (OFC)-mediated] or response-outcome (R-O) associations [presumably prelimbic cortex (PL)-mediated]. We previously found suggestive evidence that OFC and PL could compensate for each other in this task, and we more directly assessed this potential compensation here. In Experiment 1, rats received OFC, PL, combined OFC+PL, or sham lesions and then completed our devaluation task. The OFC+PL lesion group exhibited impaired devaluation. In Experiment 2, rats received cholera-toxin-b (CTb) into OFC and either neurotoxic or sham PL lesions. Rats were then sacrificed on the last training day to double-label for Arc and CTb. We found increased Arc+CTb in mediodorsal thalamus (MD) and increased Arc+ neurons in OFC when PL was lesioned, suggesting that PL lesions lead to a compensatory increased activation of the MD→OFC circuit. Our results suggest that our devaluation task can model neural compensation between OFC and PL and this compensation may be regulated by MD.

**Significance Statement:** To detect compensatory responses, behavioral models that use different strategies must be developed to determine if the strategies shift when a brain area or circuit is incapacitated. Neural compensation is commonly observed in human research but only a few models of neural compensation exist, and few identify compensation within the prefrontal cortex. This research is among the first to show neural compensation between prefrontal cortex regions and implicate a thalamocortical circuit in modulating this compensation. Not only will this model provide a way to behaviorally identify subtle neurological shifts, it can also elucidate basic neurological mechanisms that mediate how circuits interact with each other and how dysfunction in one circuit can affect connectivity in other brain areas.

## Introduction

Prodromal symptomatology, or symptoms that are below clinical threshold but signal the beginning of a disease, often appear in neuropsychiatric disorders including schizophrenia (SCZ) (Chung & Cannon, 2015; Conroy et al., 2018). Despite the neuronal dysfunction that can be tracked through neuroimaging methods, it can be hard to detect reliable behavioral changes in populations with prodromal symptomology (Barbour et al., 2010; Keshavan et al., 2002; Yaakub et al., 2013). One possible reason for unreliable or subtle behavioral changes is neural compensation, here defined as when brain regions not normally necessary to support learning for the task are recruited to utilize alternative strategies.

We intend to examine the possibility of compensatory responses in the devaluation task, as people with chronic SCZ are impaired in this task (Morris et al., 2018; Morris et al., 2015). In devaluation tasks, a subject learns that a response or a cue predicts a reward. The value of the reward is then typically reduced using motivational (e.g. selective satiety) or associative (e.g. conditioned taste aversion) methods. Devaluation is assessed by presenting the response option or cue that previously led to the devalued outcome and responding for that response option or cue is typically reduced in a test session. Traditional devaluation tasks typically seek to isolate one behavioral strategy by having either a response or a cue predict reward. This approach is valuable for isolating the circuitry of one particular strategy, but this reductionist approach is likely insensitive to detecting neurological compensation that could occur if there was more than one strategy possible and individuals could switch to a different strategy subserved by a different brain area/circuit.

To address this gap in knowledge, we developed an innovative rodent devaluation task (Fisher et al., 2017; Fisher et al., 2020; Pickens et al., 2017) that can be solved using two dissociable strategies. In our task, responses to two lever-cuelight compounds earn different rewards, and rats could guide behavior based on the cuelight (a stimulus-outcome [S-O] association), which putatively relies on lateral orbitofrontal cortex (OFC) (Gallagher et al., 1999; Panayi & Killcross, 2018; Rhodes & Murray, 2013; Rudebeck & Murray, 2011). Alternatively, the rats could guide behavior based on the lever-location (a response-outcome [R-O] association]), which putatively relies on prelimbic cortex (PL) for acquisition (Balleine & Dickinson, 1998; Corbit & Balleine, 2003; Coutureau & Killcross, 2003; Coutureau et al., 2009; Ostlund & Balleine, 2005; Tran-Tu-Yen et al., 2009) but does not rely on OFC (Ostlund & Balleine, 2007; Panayi & Killcross, 2018).

Previously, we found that inactivation of OFC or PL during operant training did not affect future devaluation, despite rats naturally preferring a R-O task strategy (Fisher et al., 2020). As previous research suggests that either OFC or PL would be needed during training for later devaluation (depending on the strategy used to guide behavior), we suggested that rats may be able to switch between strategies depending on the cortical substrates that are active during training to guide future goal-directed action (Fisher et al., 2020). Furthermore, we propose that the mediodorsal thalamus (MD) modulates PFC connectivity such that, when PL or OFC activity is disrupted, MD directs attentional resources to the brain region that remains active to encode the information needed for future goal-directed action. MD is a higher-order thalamic nucleus that is reciprocally connected to PL and OFC (Alcaraz et al., 2016; Groenewegen, 1988) and is proposed to relay salience information to modulate attention and aid in task engagement (Rikhye et al., 2018; Schmitt et al., 2017; Seeley et al., 2007).

Here, we test whether there is evidence that PL and/or OFC can guide devaluation under normal circumstances in our task (by lesioning both brain areas) and whether lesioning PL would lead to a compensatory increase in OFC neuronal activity and in the MD neurons that project to OFC.

## Methods

### Subjects

Male and female naïve Long Evans rats (n = 79) bred in-house from parents from Charles River Laboratories (Raleigh, NC and Kingston, NY) were used for all experiments. Previous research in our lab found no sex differences in devaluation (Fisher et al., 2020), so we used mixed-sex groups for Experiments 1 and 2. All animals were individually housed after surgery and maintained on a 12-hour reverse light-dark cycle (lights off at 7:00 AM) in a temperature- and humidity-controlled room.

The animals were free fed until they reached the surgery weight threshold (295 g for males and 215 g for females) and then received surgery. They remained on free feed for at least three days following surgery. They were then food-restricted to 85% of their initial free-feeding weight by daily feedings with a minimum of 5 g of food chow per day. Once rats reached their 85% target weights, the target body weight increased by 1 g/day for males or 0.25 g/day for females for the remainder of the experiment, such that the rats’ target weights gradually increased by 7 g/week or 1.75 g/week. Water was available *ad libitum*. All procedures and animal care were in accordance with the Kansas State University Institutional Animal Care and Use Committee guidelines, the National Institutes of Health Guidelines for the Care and Use of Laboratory Animals, and United States federal law.

### Behavioral apparatus

Experiments were conducted in 12 standard self-administration chambers (Med Associates, St. Albans, VT). Each chamber is equipped with a pellet dispenser that delivers a 45-mg precision pellet (catalog #1811155; TestDiet, Richmond, IN), a 45-mg chocolate-flavored sucrose pellet (catalog #1811256; TestDiet, Richmond, IN), or a 45-mg grain pellet (catalog #1812127; TestDiet, Richmond, IN). The identity of the pellet dispensed was dependent on the task phase. The chambers have two retractable levers on either side of the food cup at approximately one third of the total height of the chamber, with a white cuelight located above each lever. A red houselight is mounted on the top-center of the back wall. A speaker for delivering auditory stimuli is located on the left side of the back wall of the chambers, on the opposite wall from the food cup. A Dell Optiplex computer, equipped with Med-PC for Windows, controls the equipment and records lever-presses.

### General surgical methods

Rats underwent surgery for all experiments. Animals were anesthetized with 3-5% isoflurane in an induction chamber and then placed in a stereotaxic device (Kopf Instruments, Tujunga, CA) and maintained on 1-3% isoflurane for the duration of the surgery. Once the skull was exposed, bregma was used to determine injection placement, for which holes were burred with a drill. All infusions (described below for each individual experiment) were at a rate of 0.1 μl/min. Following the infusion, the needle was left in place for 5-7 min to allow for diffusion. After the needle was removed, the incision was closed with wound clips and triple antibiotic ointment was applied. Animals were given 1 mg/kg flunixin (Norbrook, Overland Park, KS) subcutaneously upon completion of surgery and 24-h later during the post-op health check.

### Devaluation task

#### Training (Experiments 1-2)

After at least a 10-day recovery period, rats received 3 once-daily 40 min magazine training sessions, one with each pellet type, with pellets being delivered every 120 sec. Then, they received 4 lever press training sessions (2 with each lever; twice daily) on a fixed-interval-1 schedule earning grain pellets with no cuelights present. Here, lever presses could earn a reward with a press every one second. Sessions ended when 50 rewards were earned or 60 min passed, whichever came first. Grain pellets and no cuelights were used to prevent the formation of R-O and S-O associations with the pellets used as reinforcers in upcoming cued-trial operant training. Next the rats received 4 once-daily cued-trial operant training sessions, two sessions for each cuelight-lever-reinforcer combination. Only one cuelight-lever-pellet compound was available during a session. Throughout training, left-lever presses earned precision pellets and were always associated with illumination of a steady cuelight (the white cuelight above the left lever). Right-lever presses earned chocolate pellets and were always associated with flashing illumination of a cuelight (the white cuelight above the right lever). Every cued-trial training session was 40 min long with 40 trials. All trials were 40 sec long and rats could earn up to two pellets per trial for lever-pressing, one in the first half and one in the second half of the trial at randomized times. During the inter-trial interval, the levers retracted and the cuelights turned off.

#### Devaluation Choice Test (Experiment 1 only)

Following cued trial training, a choice test was administered. In the operant chambers, rats received a 1-hr satiation session with access to 30g of either chocolate or precision pellets (counterbalanced) presented in ceramic bowls. Fifteen minutes after the satiation period, rats received a 12-min choice test with twelve 40-sec trials. During each trial, both levers with the associated cuelights were presented and responding was measured. The cuelights above the levers were in the same position as during training (e.g.: flashing cuelight above the left lever). Responding should be decreased for the cuelight-lever compound that earns the devalued reinforcer if rats devalue properly. Lever presses did not earn pellets to ensure the rats used their memory representations of the cue-lever combinations to guide behavior. Next, rats received two cued-trial retraining sessions, one with each levercuelight-food compound, to return lever pressing rates to baseline. Rats then completed another choice test with the opposite pellet devalued during the satiation period (**Fig. 1A**).

**Figure 1.**
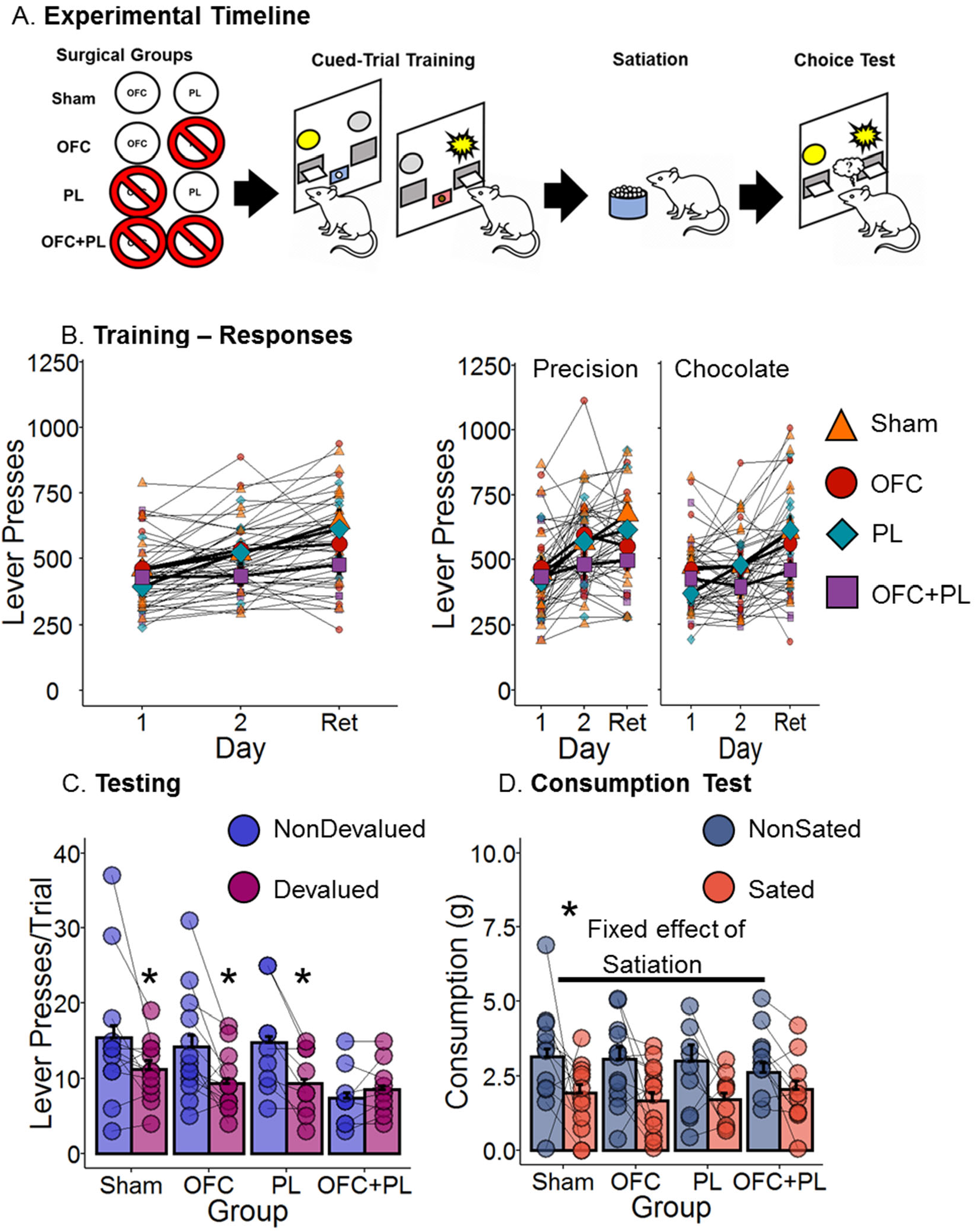
Experiment 1 Timeline and Experimental Data. **A.** Schematic of our surgical and devaluation task procedure for Exp. 1. Red circle with the slash represents the region that was lesioned. Blue background on the foodcup represents precision pellets. Pink background on the foodcup represents chocolate pellets. Blue bowls represent sated on precision pellets. **B.** Lever presses/session (mean ± SEM) in each lesion group during the first and second training cued-trial training days and retraining days for each pellet in Exp. 1. Left panel is averaged across and pellet type and right panel is separated by pellet type. Orange triangles = Sham group. Red circles = OFC group. Teal diamonds = PL group. Purple squares = OFC+PL group. **C.** Lever presses/Trial (mean ± SEM) in each lesion group during the devaluation te st in Exp. 1 (averaged across trial and test). Blue bars represent responding on the nondevalued lever. Maroon bars represent responding on the devalued lever. **D.** Consumption in grams (mean ± SEM) of the sated and nonsated pellet types during the consumption test in Exp. 1. Grayish blue bars = nonsated pellet type. Brownish orange bars = sated pellet type. *=p<0.05

#### Devaluation Consumption Test (Experiment 1 only)

Finally, at least one day after the final choice test, animals in Experiment 1 completed 2 consumption tests. Each consumption test included a 60-min satiation period with one of the two pellet types (counterbalanced across group), identical to the satiation periods that preceded the choice tests. Fifteen minutes following the end of the satiation period, the animals had access to two ceramic bowls in the operant chambers. One bowl contained 10g of precision pellets and the other bowl contained 10g chocolate pellets. The consumption test lasted 10 min. After completion of the test, the remaining pellets were sorted by type and weighed to determine consumption. Animals received at least one day off after the first consumption test before completing the second consumption test identical to the first, except that animals were sated on the opposite pellet type as in the first consumption test.

### Individual Experimental Designs

#### Experiment 1

Male and female rats (n = 59; 12-20/group) received neurotoxic lesions and/or sham lesions of PL and OFC (**Fig. 1A**). There were four surgical groups: PL (13 rats; 6 male, 7 female), OFC (14 rats; 7 male, 7 female), OFC+PL (20 rats; 10 males, 10 females), and Sham (12 rats; 6 males, 6 females). The PL and OFC groups received a bilateral lesion of PL or OFC, respectively. The OFC+PL group received two bilateral lesions: a bilateral OFC lesion and a bilateral PL lesion. All excitotoxic lesions were created by infusing 0.4 μl (OFC) of 20 mg/ml NMDA and/or 0.3 μl (PL) of 10 mg/ml NMDA in phosphate-buffered saline (PBS). The Sham group received infusions of PBS to OFC and PL. Single lesion groups also received an infusion of PBS (0.4 μl or 0.3 μl) into the non-lesioned area. The OFC infusion was at the following coordinates relative to bregma: anterior-posterior (AP): +3.5 mm, medial-lateral (ML): +3.3 mm, dorsal-ventral (DV): −5.4 mm. The PL infusion was at the following coordinates relative to bregma: AP: +3.2 mm, ML: +0.7 mm, DV: −3.6 mm. Following surgery, the rats began cued-trial training, and then completed the Cue Normal devaluation choice tests and devaluation consumption tests as described above.

#### Experiment 2

Male and female rats (n = 20; 8-12/group) received neurotoxic lesions or sham lesions of PL (**Fig. 2A**). Rats in the PL lesion group (12 rats; 6 males, 6 females) received bilateral infusions of 0.3 μl of 10 mg/ml NMDA. Rats in the PL sham group (8 rats; 3 males, 5 females) received infusions of 0.3 μl of PBS. The PL infusions were at the following coordinates relative to bregma: AP: +3.2 mm, ML: +0.7 mm, DV: −3.6 mm. In addition, all rats in both groups received a unilateral, side counter-balanced, 0.2 μl infusion of 1% CTb conjugated with Alexa Fluor 488 (Invitrogen) in PBS in OFC. The OFC infusion was at the following coordinates relative to bregma: AP: +3.5 mm, ML: +3.3 mm, DV: −5.4 mm. Following surgery, the rats began cued-trial training at least 3 weeks later to allow for retrograde transport and were perfused 75 min following the start of the last session of training (the 4^th^ session) to optimize Arc protein expression (Lonergan et al., 2010).

**Figure 2.**
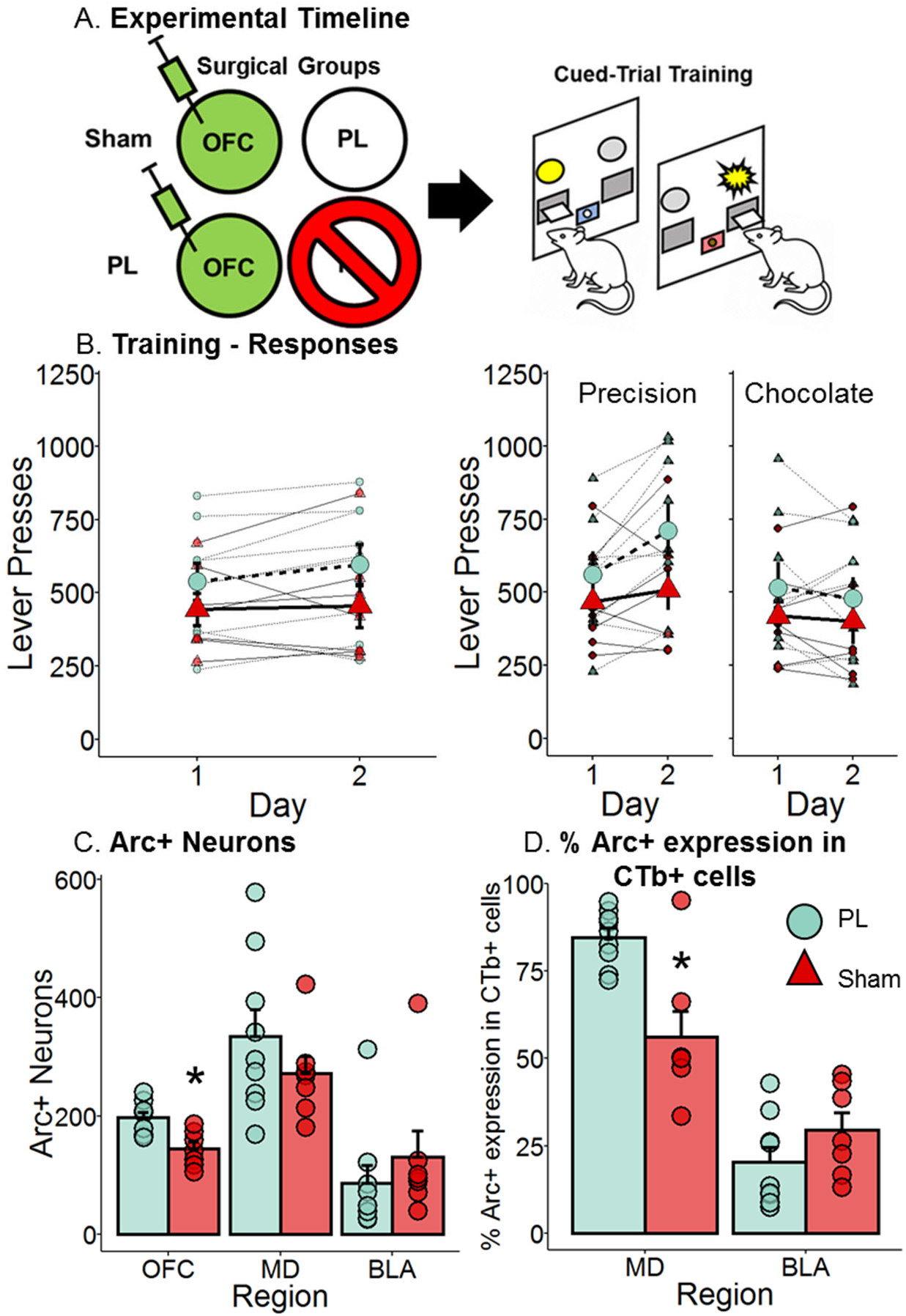
Experiment 2 Timeline and Experimental Data. **A.** Schematic of our surgical and devaluation task procedure for Exp. 1. Green syringe represents an infusion of cholera-toxin b. Red circle with the slash represents the region that was lesioned. Blue background on the foodcup represents precision pellets. Pink background on the foodcup represents chocolate pellets. **B.** Lever presses/session (mean ± SEM) in each lesion group during the first and second training cued-trial training days and retraining days for each pellet in Exp. 1. Left panel is averaged across and pellet type and right panel is separated by pellet type. Aqua circle = PL group. Red triangle = Sham group. **C.** Arc+ neurons (mean ± SEM) in the OFC, MD, and BLA. Aqua bar= PL group. Red bar = Sham group. **D.** Percent Arc+ expression in CTb+ cells in the MD and BLA. Aqua bar= PL group. Red bar = Sham group. *=p<0.05

### Histological Procedures

#### Perfusions

75 minutes following the beginning of the last session of training (Experiment 2) or after the completion of behavioral testing (Experiment 1), the rats were deeply anesthetized with pentobarbital and perfused with 100 ml of 0.1-M PBS followed by 400-500 ml of 4% paraformaldehyde (PFA). The brains were removed, post-fixed in PFA for 2 hr, and stored in 0.1-M PBS with 30% (w/v) sucrose for 48-72 hr until dehydrated. Sections (40-μm) were taken from each brain using a cryostat, and every section was collected.

#### Experiment 1-2

Every 3^rd^ (Exp. 1) or 4^th^ (Exp. 2) brain section was mounted and Nissl stained using thionin. Lesions were verified using a light microscope (model BX41, Olympus) and SPOT 5.1 Advanced Software. Only rats with bilateral lesions with 60% or more damage of the target area on a section were included in the data analyses.

#### Experiment 2

Immunohistochemistry (IHC) using an immunofluorescence method was performed to label for Arc and CTb (Experiment 2) by adapting the IHC procedure used by Marchant and colleagues (Marchant et al., 2010).

The tissue was gently agitated on a plate shaker throughout. First, the free-floating tissue was washed repeatedly with 0.1-M PBS (pH 7.4). The sections were blocked for 1 hr with 10% normal horse serum (NHS), 0.5% Triton-X, and 5% bovine serum albumin (BSA) in PBS and incubated at room temperature for 48 hr in rabbit anti-Arg 3.1 (1:4,000; Cell Signaling) and goat anti-CTb (1:5,000; ListLabs) diluted with 10% NHS, 0.5% Triton-X, 5% BSA, and 0.1% sodium azide in PBS. After three PBS rinses, sections incubated for 6 hr at room temperature in donkey anti-rabbit Alexa Fluor 594 (1:200) and donkey anti-goat Alexa Fluor 488 (1:200) diluted with 10% NHS, 0.5% Triton-X, and 5% BSA in PBS. The tissue was rinsed in PBS and incubated for 10 min at room temperature in DAPI (1:25,000) in PBS. The tissue was rinsed twice for 1 min each and then stored in PBS at 4°C until they were mounted on charged slides and coverslipped with mounting medium (90% glycerol and 0.5% n-propyl gallate in 20mM Tris, pH 8.0). Uniform stitched images (Zeiss 700) at 40X magnification for each area at the following coordinates, relative to bregma, were used for the analyses: OFC (+4.2), MD (−2.9), and the basolateral amygdala (BLA) (−3.0).

### Statistical analysis

For the behavioral data, multilevel Poisson regressions were run with a negative binomial distribution on the number of lever presses (Training and Testing) or a Poisson distribution on the number of rewards earned (Training). The best fitting models were chosen via model comparisons that employed the Akaike Information Criterion (AIC), which took into account the number of parameters entered into the model and penalized more complex models (Kuha, 2004). For all experiments, the best fitting models were a full factorial analysis. For neurological analyses, the data were analyzed using two-tailed Student’s *t*-tests with the dependent variables including the number of Arc+ neurons in OFC, BLA, and MD or total number of CTb+Arc neurons divided by the total number of CTb+ neurons in MD and BLA. As with our lesions, we define OFC as lateral OFC for immunohistochemistry assays. All categorical predictors were effect coded. For all analyses (including t-tests), the residual distributions were examined for normality homoscedasticity, and linearity and any violations and subsequent transformations are reported in the appropriate results section below.

#### Experiment 1

For the training data, possible predictors included the between-subjects variables PL (Sham, PL) and OFC (Sham, OFC) and the within-subjects variables Training Day (Day 1, Day 2, Retrain) and Pellet Type (Precision, Chocolate). The random effects structure included Intercept (subject) and two slopes (Training Day and Pellet Type). For the test data, possible predictors included the between-subjects variables PL (Sham, PL) and OFC (Sham, OFC) and the within-subjects variables Lever (Nondev, Dev) and Trial. The random effects structure included Intercept (Subject) and two slopes (Lever and Trial). Planned comparisons were run for significant interactions including PL and OFC using Type I Sums of Squares.

#### Experiment 2

The training data included the possible predictors: PL (Sham, PL), Training Day (Day 1, Day 2, Retrain), and Pellet Type (Precision, Chocolate). The random effects structure included Intercept (subject) and two slopes (Training Day, and Pellet Type). Planned comparisons were run for significant interactions including PL using Type I Sums of Squares. The neurological data were analyzed with a Student’s independent samples *t*-test with the variable Group (PL, Sham) on the ratio of double-labeled CTb+Arc neurons to the number of CTb+ neurons in MD and BLA or on the number of Arc+ neurons in each MD, BLA, and OFC.

All data were analyzed with R (R 3. 3. 1). Significant interactions (*p* < 0.05) of the highest order with the variables PL and OFC (Exp. 1) or PL (Exp. 2) were followed by planned comparisons using emmeans.

## Results

### Experiment 1

#### Histological exclusions

For both PL and OFC, lesions had to damage at least 60% of the region on a section. Based on these criteria, 15 rats (1 Male-OFC, 2 Male-PL, 2 Female-PL, 5 Male-OFC+PL, 5 Female-OFC+PL) were excluded. The final group sizes in Experiment 1 were as follows: OFC, n = 13 (6 Males, 7 Females); PL, n = 9 (4 Males, 5 Females); OFC+PL, n = 10 (5 Males, 5 Females); and Sham, n = 12 (6 Males, 6 Females).

#### Training

The number of responses across training days did not interact with PL and OFC groups (**Fig. 1B**). A multilevel Poisson regression with a negative binomial distribution was used to analyze the data. Comparing AICs, the three tested models were all within 5 points of each other. Therefore, we decided to use the full factorial model which included OFC × PL × Pellet × Day and all lower effects. Intercept (Subject) and the slopes of Pellet and Day were added as random effects. There was significant OFC × Day interaction (*b* = −0.06, *z* = −2.89, *p* < 0.01), but as this effect did not also include the PL group, no comparisons were run. Other lower-level effects are shown in Table 1.

**Table 1 –.** Experiment 1 Training: Responses.

|  | *b* | *z*-value | *p* |
| --- | --- | --- | --- |
| Intercept | 6.19 | 160.37 | **<0.01** |
| PL | -0.06 | -2.35 | **0.02** |
| OFC | 0.03 | 1.05 | 0.29 |
| Pellet | -0.05 | -2.40 | **0.02** |
| Day | 0.13 | 6.47 | **<0.01** |
| PL*OFC | 0.01 | 0.36 | 0.72 |
| PL*Pellet | -0.002 | -0.12 | 0.90 |
| OFC*Pellet | 0.01 | 0.33 | 0.75 |
| PL*Day | 0.01 | 0.25 | 0.80 |
| OFC*Day | -0.06 | -2.89 | **<0.01** |
| Pellet*Day | -0.01 | -0.40 | 0.69 |
| PL*OFC*Pellet | 0.01 | 0.41 | 0.68 |
| PL*OFC*Day | -0.02 | -0.97 | 0.33 |
| PL*Pellet*Day | 0.01 | 0.68 | 0.50 |
| OFC*Pellet*Day | 0.001 | 0.07 | 0.94 |
| PL*OFC*Pellet*Day | -0.02 | -1.24 | 0.22 |

Note: Data are effect coded. Bolded value are significant ( $p < .05$ )

The number of rewards earned increased across the training days but there were no group differences in this effect (data not shown). A multilevel Poisson regression with a Poisson distribution was used to analyze the data. We used the same full factorial model structure as that used for the number of responses which included OFC × PL × Pellet × Day and all lower effects. Intercept (Subject) and the slopes of Pellet, and Day were added as random effects. No with effects or interactions with OFC and PL were significant (all *p* > .05; Table 2).

**Table 2 –.** Experiment 1 Training: Rewards Earned.

|  | *b* | *z*-value | *p* |
| --- | --- | --- | --- |
| Intercept | 4.28 | 551.16 | **<0.01** |
| PL | 0.002 | 0.33 | 0.74 |
| OFC | 0.007 | 0.98 | 0.33 |
| Pellet | -0.004 | -0.54 | 0.59 |
| Day | 0.04 | 5.00 | **<0.01** |
| PL*OFC | -0.004 | -0.53 | 0.59 |
| PL*Pellet | 0.008 | 1.04 | 0.30 |
| OFC*Pellet | -0.0005 | -0.07 | 0.95 |
| PL*Day | 0.01 | 1.25 | 0.21 |
| OFC*Day | -0.01 | -1.25 | 0.21 |
| Pellet*Day | -0.01 | -1.16 | 0.25 |
| PL*OFC*Pellet | -0.004 | -0.48 | 0.63 |
| PL*OFC*Day | -0.0003 | -0.04 | 0.97 |
| PL*Pellet*Day | -0.003 | -0.37 | 0.71 |
| OFC*Pellet*Day | -0.001 | -0.13 | 0.90 |
| PL*OFC*Pellet*Day | -0.01 | -1.09 | 0.28 |

Note: Data are effect coded. Bolded value are significant ( $p < .05$ )

#### Devaluation test

The Sham, OFC, and PL groups made more responses on the nondevalued lever compared to the devalued lever, indicative of a devaluation effect, but the OFC+PL group did not (**Fig. 1C**). A multilevel Poisson regression with a negative binomial distribution was used to analyze the data. Comparing AICs, the best fitting model (AIC = 3579.8) was about 8 points better than the next best model (AIC = 3543.4). The best fitting model included OFC × PL × Lever × Trial and all lower effects. Intercept (Subject) and the slopes of Lever and Trial were added as random effects. There was a significant OFC × PL × Lever interaction (*b* = −0.09, *z* = −2.72, *p* < 0.01). Other lower-level effects are shown in Table 3. The OFC × PL × Lever interaction revealed that the Sham (*z* = 4.70, *p* < 0.01), OFC (*z* = 5.54, *p* < 0.01), and PL (*z* = 3.48, *p* < 0.01) groups responded on the nondevalued lever significantly more than the devalued lever while the OFC+PL group (*z* = −1.59, *p* = 0.11) did not.

**Table 3 –.** Experiment 1 Devaluation Test.

|  | *b* | z-value | *p* |
| --- | --- | --- | --- |
| Intercept | 2.29 | 29.99 | **<0.01** |
| PL | -0.28 | -5.35 | **<0.01** |
| OFC | -0.07 | -1.35 | 0.18 |
| Lever | 0.13 | 3.88 | **<0.01** |
| Trial | -0.15 | -5.18 | **<0.01** |
| PL*OFC | 0.02 | 0.37 | 0.72 |
| PL*Lever | -0.07 | -2.00 | 0.045 |
| OFC*Lever | -0.05 | -1.63 | 0.10 |
| PL*Trial | -0.11 | -4.54 | **<0.01** |
| OFC*Trial | 0.01 | 0.37 | 0.71 |
| Lever*Trial | 0.001 | 0.07 | 0.94 |
| PL*OFC*Lever | -0.09 | -2.72 | **<0.01** |
| PL*OFC*Trial | 0.03 | 1.55 | 0.12 |
| PL*Lever*Trial | -0.01 | -0.52 | 0.60 |
| OFC*Lever*Trial | 0.02 | 1.08 | 0.28 |
| PL*OFC*Lever*Trial | 0.001 | 0.05 | 0.96 |

Note: Data are effect coded. Bolded value are significant ( $p < .05$ )

#### Consumption test

The rats ate significantly more of the pellet type they were not sated on (i.e. the nondevalued pellet type) regardless of group (**Fig. 1D**). A multilevel Gaussian regression was used to analyze the data. The model structure included OFC × PL × Satiation and all lower effects. Intercept (Subject) and the slope of Satiation were added as random effects. There was a significant effect of Satiation (*b* = 0.55, *t* = 4.01, *p* < 0.01). There was no significant interaction of OFC × PL × Satiation (*b* = −0.11, *t* = −0.81, *p* = 0.42). No other effects or interactions were significant (*p* > 0.05; Table 4). The rats ate more of the nondevalued pellet type (*M* = 2.97, *SE* = 0.21) compared to the devalued pellet type (*M* = 1.83, *SE* = 0.16).

**Table 4 –.** Experiment 1 Consumption Test.

|  | *b* | t-value | *p* |
| --- | --- | --- | --- |
| Intercept | 2.40 | 17.21 | **<0.01** |
| OFC | -0.05 | -0.36 | 0.72 |
| PL | -0.05 | -0.38 | 0.71 |
| Satiation | 0.56 | 4.01 | **<0.01** |
| OFC*PL | 0.04 | 0.26 | 0.79 |
| OFC*Satiation | -0.07 | -0.49 | 0.63 |
| PL*Satiation | -0.09 | -0.68 | 0.50 |
| OFC*PL*Satiation | -0.11 | -0.81 | 0.42 |

Note: Data are effect coded. Bolded value are significant ( $p < .05$ )

### Experiment 2

#### Histological exclusions

For PL, lesions had to damage at least 60% of the region on a section. Based on these criteria, 4 rats (1 Female, 3 Males) were excluded. In addition, 1 Sham Male was excluded due to extensive cortical damage that was not the result of the brain extraction. The final group sizes in Experiment 1 were as follows: PL, n = 9 (4 Males, 5 Females); Sham, n = 7 (2 Males, 5 Females).

#### Training

PL lesions did not affect responding during training **(Fig. 2B**). A multilevel Poisson regression with a negative binomial distribution was used to analyze the data with the same structure as Experiment 1, which included PL × Pellet × Day and all lower effects. Intercept (Subject) and the slopes of Pellet and Day were added as random effects. No effects including PL were significant. Other effects are shown in Table 5.

**Table 5 –.** Experiment 2 Training Responses.

|  | *b* | z-value | *p* |
| --- | --- | --- | --- |
| Intercept | 6.14 | 69.24 | **<0.01** |
| PL | 0.11 | 1.28 | 0.2 |
| Pellet | -0.11 | -3.07 | **<0.01** |
| Day | 0.03 | 0.67 | 0.5 |
| PL*Pellet | -0.02 | -0.42 | 0.67 |
| PL*Day | 0.04 | 0.81 | 0.41 |
| Pellet*Day | -0.12 | -3.28 | **<0.01** |
| PL*Pellet*Day | -0.04 | -1.14 | 0.25 |

Note: Data are effect coded. Bolded value are significant ( $p < .05$ )

The number of rewards earned did not differ across any variable (data not shown). A multilevel Poisson regression was used to analyze the data. When we examined model fits for the model analyzing Rewards Earned, the PL × Pellet × Day and all lower effects model structure had a substantially lower AIC than the PL × Pellet, Pellet × Day, and all lower effects model. We used the same model structure as that used for the number of responses which included PL × Pellet × Day and all lower effects. Intercept (Subject) and the slopes of Pellet, and Day were added as random effects. There were no significant effects or interactions of any variable (all *p* > .05; Table 6).

**Table 6 –.** Experiment 2 Training Rewards Earned.

|  | *b* | *z*-value | *p* |
| --- | --- | --- | --- |
| Intercept | 4.25 | 199.74 | **<0.01** |
| PL | 0.006 | 0.30 | 0.76 |
| Pellet | -0.03 | -1.68 | 0.09 |
| Day | 0.01 | 0.42 | 0.68 |
| PL*Pellet | -0.006 | -0.34 | 0.73 |
| PL*Day | 0.02 | 0.69 | 0.49 |
| Pellet*Day | -0.01 | -0.42 | 0.67 |
| PL*Pellet*Day | -0.01 | -0.48 | 0.63 |

Note: Data are effect coded. Bolded value are significant ( $p < .05$ )

#### Neurological analyses

The residual distribution of the Arc+ neurons in BLA were not normal. To correct for non-normality, the number of Arc+ neurons in BLA were log transformed prior to analysis. There was an influential point in the PL group for the ratio of Arc and CTb neurons in MD that created a slight skew (see highest PL point in **Fig. 2C**). The analysis was run with and without the data point and did not alter the results. Therefore, we decided to include the data point.

Student’s independent two samples *t*-tests were used to analyze the overall number of Arc+ neurons in MD, OFC, and BLA (**Fig. 2C**) or the percentage of CTb+ neurons that were also Arc-positive in MD and BLA (**Fig. 2D**). While the numbers of Arc+ cells in MD, *t*(14) = 1.11, *p* = .28, and BLA, *t*(14) = −.84, *p* = .41, were not different between the PL lesioned rats compared to Sham rats, there were more Arc+ neurons in the OFC in PL lesioned rats compared to Sham rats, *t*(14) = 3.74, *p* < .01 (Fig. 2C). While in the BLA the percentage of CTb neurons that were Arc+ were not different between PL lesioned and Sham rats, *t*(14) = −1.42, *p* = .18, there was a higher percentage of CTb neurons that were Arc+ in the MD in the PL lesioned group compared to the Sham group, *t*(14) = 3.98, *p* < .01.

## Discussion

In the current experiments, we showed that rats can exhibit normal goal-directed action in our task when either PL or OFC is functional but goal-directed action is impaired when both PL and OFC are damaged. We also showed that when PL is lesioned, neuronal activity in both OFC and MD->OFC projecting neurons is increased during training. Below, we discuss how the effect of OFC+PL lesions on devaluation suggests our task can model neural compensation. We also discuss the increased neural activity in OFC and MD during training and the role of MD in modulating neural compensation.

### OFC+PL lesions impaired devaluation

Our behavioral results suggest that least one of these regions (PL or OFC) is necessary and sufficient for training and/or performance in our devaluation task with congruent cue-compounds during training and testing. Our task is designed in such a way that it can be solved using an S-O strategy by paying attention to the unique cuelights above the levers, likely mediated by OFC (Gallagher et al., 1999; Panayi & Killcross, 2018; Rhodes & Murray, 2013; Rudebeck & Murray, 2011), or an R-O strategy by paying attention to the spatial lever location, likely mediated by PL during the operant training (Balleine & Dickinson, 1998; Corbit & Balleine, 2003; Coutureau & Killcross, 2003; Coutureau et al., 2009; Ostlund & Balleine, 2005; Tran-Tu-Yen et al., 2009). Previously, we have shown that OFC or PL inactivation during training does not impair future devaluation in our task, despite rats preferring to use an R-O strategy (Fisher et al., 2020). In Exp. 1 in the current paper, we found similar results with neurotoxic lesions of either of the regions, as long as the other area was intact. This suggests that, while the R-O strategy was the default strategy learned, the cue-based strategy could be learned and used to express normal goal-directed action if PL was exhibiting dysfunction during training. Our current results support our hypothesis that compensation between PL and OFC can maintain goal-directed action as long as the cuelights and lever-location predict the same outcome during the test. Lesioning *both* areas (preventing any compensation between them) disrupted the pattern of goal-directed action observed.

Our findings do not invalidate other devaluation procedures where S-O (e.g., Pavlovian) or R-O (e.g., free-operant) strategies are used exclusively to guide behavior. Indeed, our interpretation of our data is based heavily on results of these tasks showing a dissociation between tasks that are OFC or PL dependent. However, our findings and task may be more representative of how devaluation occurs in humans with multiple sources of information available to guide behavior. Our task represents one possible way to model devaluation in complex environments while leveraging the control gained from use of animal models. Therefore, our task represents a devaluation task variant that models more complex environments that may, in turn, better model goal-directed action and the underlying neural substrates in humans, contrasted with other (very useful and highly controlled) variants used to isolate the role of specific types of associations.

### Neural activity increased in OFC but not MD or BLA during training when PL was lesioned

Analysis of the Arc+ neurons in OFC, BLA, and MD found increased numbers of Arc+ neurons in OFC in PL lesioned rats and no differences between groups in MD or BLA during training. Broadly, these findings suggest that increased neural activity in OFC may represent increased learning of the S-O strategy to guide future behavior in the absence of PL competition. These findings also suggest that the population of MD neurons active during training may be altered after PL lesions (see the next section) but overall MD or BLA activity does not need to be increased for compensation to occur.

Although the increased Arc+ neurons in OFC in the PL group suggests that OFC becomes more active so that the S-O strategy may be learned in order to guide behavior when PL is damaged, we cannot say for certain that the increase in OFC activity would result in the S-O strategy being used to guide behavior. Future studies could use methods, such as exciting OFC during training, to determine whether rats devalue using the R-O or S-O strategy when OFC activity is increased during training.

### MD->OFC, but not BLA->OFC, neural activity increased during training when PL was lesioned

We found a higher ratio of Arc+CTb+ neurons in MD when PL is lesioned, revealing that MD->OFC projecting neurons are more active when PL is lesioned prior to training. This is indirect evidence for a process in which S-O and R-O strategies compete for attentional resources. Based on our data, the R-O strategy is more salient, as it is the preferred strategy used when the two strategies are pitted against each other in intact rats. If so, then the R-O association may overshadow the S-O association (or in terms of environmental stimuli, the spatial location of the lever overshadows the light) (Fisher et al., 2020). Correspondingly, OFC receives less excitatory input from MD when PL is intact, as a potential neurobiological correlate to this overshadowing/competition. However, when PL is inactive or damaged, there is no competition from the R-O association/lever spatial location and attentional resources may be shifted to OFC so the S-O/cuelight strategy can be learned. As such, our finding that MD->OFC neurons are more active when PL is damaged provides indirect but compelling evidence that MD directs attentional resources to OFC so that a cuelight strategy is learned preferentially when PL is damaged, in line with its proposed role in relaying salience information to modulate attention and aid in task engagement (Rikhye et al., 2018; Schmitt et al., 2017; Seeley et al., 2007).

Notably, we found no differences in BLA activity or BLA->OFC activity, suggesting that BLA is not involved in directly modulating compensation between OFC and PL in our task. This lack of difference was interesting given the documented role of BLA connections with OFC affecting S-O dependent tasks (Lichtenberg et al., 2017; Sias et al., 2021). While the direct BLA->OFC connections may not be involved, this finding does not rule out BLA involvement as BLA may modulate MD, influencing activity of thalamocortical projections. More research is needed to elucidate BLA’s exact role or lack thereof in neural compensation during our task.

### Conclusions

Our results indicate that our multi-response/multi-reinforcer devaluation task with cued trials can successfully model neural compensation between OFC and PL. These results provide a strong basis for future research examining the mechanisms of how OFC and PL actively compensate for each other in our task. We also found indirect evidence for our overall hypothesis that MD regulates neural compensation and attentional activity based on the salience of the strategies available and cortical availability. Further, this suggests that MD dynamically regulates attentional control of complex learning strategies between PFC brain regions. Future studies should provide a causal link that MD modulates neural compensation and determine whether this compensation may be occurring in disorders/models of disorders that affect function of PL, OFC, or both, including in prodromal SCZ.

## Acknowledgements

We would like to thank Dr. Michael Young for statistical advice and the Confocal Core supported by CVM-KSU. This work was supported by the National Institutes of Health [grant number P20 GM13109-01A1 (C.L.P)], Psi Chi [Graduate Research Grant (H.F.)], the Histochemical Society [Cornerstone Grant (H.F.)], Kansas State Graduate School [Arts, Humanities, and Social Sciences Small Grant Program (H.F.)], and Sigma Xi [Grants in Aid of Research (H.F.)].

## Notes

### Competing Interest Statement

The authors have declared no competing interest.

